# Tumors cells with mismatch repair deficiency induce hyperactivation of Pyroptosis resistant to cell membrane damage but are more sensitive to Co-treatment of IFN-γ and TNF-α to PANoptosis

**DOI:** 10.1101/2023.07.28.550878

**Authors:** Huiyan Li, Hengli Ni, Ying Li, Aijun Zhou, Xiaokang Qin, Yuqin Li, Liheng Che, Hui Mo, Chao Qin, Jianming Li

## Abstract

Hypermutated neoantigens in cancers with DNA mismatch repair deficiency (dMMR) are prerequisite for favorable clinical responses to immune-checkpoint blockade (ICB) therapy. However, TMB is not significantly associated with favorable prognosis from Preclinical and clinical studies. It implys that except for TMB, other mechanisms should be needed to contribute to successful cancer immunotherapy. We found that the hyperactivation of PANoptotic effective molecules in dMMR tumor cells caused cell membrane damage, and induced ESCRT mediated membrane repair, and protectd tumor cells from the damage caused by Triton X100, while DNA mismatch repair proficient (pMMR) tumor cells were sensitive to Triton X100 mediating cell membrane damage due to the lack of ESCRT mediated membrane repair. There were hyperactivation of GSDMD, GSDME and p-MLKL in dMMR tumor cells. Co-treatment of IFN-γ and TNF-α induced rapid death of dMMR tumor cells by inducing PANoptosis including pyroptosis, apoptosis not necrosis. pMMR tumor cells had defects in PANoptosis pathway and were resistant to co-treatment of IFN-γ and TNF-α. In conclusion, we can activate immune cells to release IFN-γ and TNF-α to overcome resistance to ICB treatment.

## INTRODUCTION

The DNA mismatch repair (MMR) pathway maintains DNA-replication fidelity, and deficiency of MMR (dMMR) is prevalent in a range of cancer types (1). One of characteristics of dMMR cancers is easy to produce high tumor mutational burden (TMB). Cancers with high mutational burden are easy to induce high immune cells infiltration in themselves, accounting for high sensitivity to immune-checkpoint blockade (ICB) therapy(2, 3) and favorable prognosis. Hypermutation-generated neoantigens are prerequisite for favorable clinical responses to ICB(2–4) (5). However, TMB is not significantly associated with favorable prognosis from Preclinical and clinical studies, the objective response rate (ORR) to anti-PD-1 therapy is quite variable, ranging from 28% to 53% (3, 6); (7). These observations imply that except for TMB, there are other unknown mechanisms to clarify for successful cancer immunotherapy(8).

TMB in tumor is usually positively correlated with activation of T cell. CD8+ T cells are prerequisite for favorable successful immune-checkpoint blockade (ICB) therapy. Pre-existing CD4+ T cells are helpful for CD8 T cell activation and long survival in the tumor microenvironment, and finally are more effective for ICB therapy(9–11). And the IFN-γ and TNF-α are important cytosolic and activative molecules. Mutations in *B2M* or *JAK1/2*, key genes for IFN signal transduction in cancer cells can attenuate T cells cytotoxic attack, which is resistant for ICB therapy in dMMR cancers (12). Moreover, impairing tumor infiltration of T cells is a choice for dMMR cancers resistant to ICB therapy. Specially, some research showed that deficiency of MLH1 and subsequent accumulation of cytosolic DNA activate the cGAS-STING pathway, contributing to increased immunity(8). It means that there are many unknown mechanisms that account for that the primary dMMR tumors are resistant to ICB(13).

Successful cancer immunotherapy not only need immune system activation including more activitive T cells specially targeting cancer, dendritic cells, macrophage, B cells and so on, but also need cancer sensitive to killing moleculars (such as proforin, granzyme, TNF-α, INF-γ) releasing by T cells and NK cells, if cancer is resisitant to the killing moleculars, they can escape from killing by T cells.

Cytotoxic T lymphocytes (CTLs) and natural killer cells kill virus-infected and tumor cells through the polarized release of perforin, granzymes, IFN-γ and TNF-α. Perforin can form pores in the outside plasma membrane of the target cell to let granzymes enter the cytosol and initiate PANoptosis including pyroptosis, apoptosis and necrosis(14). While TNF-α-mediated signaling is a pro-survival signaling(15, 16), IFN-γ-mediated signaling is cytotoxic. However, these pathways have counteracting effects. TNF-α and IFN-γ together sensitize the cells to undergo PANoptosis without suppressing TNF-α inducing cell death by IFN-γ(17). If cancer get the deficiency in PANoptosis pathway, they are risisitant to the cancer immunotherapy treatment including ICB treatment.

Here, we show deficiency of MLH1 and subsequent accumulation of cytosolic DNA activate AIM2-ZBP-ASC-RIPK1-RIPK3-CASP8-CASP1-GSDMD-GSDME-MLKL pathway, contributing to increased cytotoxic reaction, inducing the dMMR cancers death.

We found that ESCRT proteins were precisely recruited in dMMR tumor cell. Inhibition of ESCRT machinery in cancer-derived cells enhanced their susceptibility to Triton X100 mediating cell killing. The hyperactivation of PANoptotic effector molecules in dMMR tumor cells caused cell membrane damage, and induced ESCRT mediated membrane repair, and protectd tumor cells from the damage caused by Triton X100, while pMMR tumor cells were sensitive to Triton X100 mediating cell membrane damage due to the lack of ESCRT mediated membrane repair. There were hyperactivation of GSDMD, GSDME and p-MLKL in dMMR tumor cells. Co-treatment of IFN-γ and TNF-α induced rapid death of dMMR tumor cells by inducing PANoptosis including pyroptosis, apoptosis not necrosis. pMMR tumor cells had defects in PANoptosis pathway and were resistant to co-treatment of IFN-γ and TNF-α.

Our findings reveal additional mechanisms of responsiveness versus unresponsiveness to ICB therapy in dMMR cancer hosts, and provide directions for future clinical practice.

## Results

### dMMR tumor cell is more sensitive than pMMR tumor cell to PANoptosis with co-treatment of IFN-*γ* and TNF-*α*

We induced dMMR tumor cell lines DLD1 or HCT116 to PANoptosis with IFN-γ, TNF-α or co-treatment of IFN-γ and TNF-α compare with pMMR tumor cell lines SW480. The result showed IFN-γ induced little death in SW480 or HCT116, except for DLD1; TNF-α could induced HCT116 more death than SW480, but lillte death in DLD1; Co-treatment of IFN-γ and TNF-α induced most cell death in DLD1, HCT116 or SW480. The result suggested that dMMR tumor cell was more sensitive than pMMR tumor cell to PANoptosis with co-treatment of IFN-γ and TNF-α (Figure 1A, 1C, 1D).

**Figure 1.**
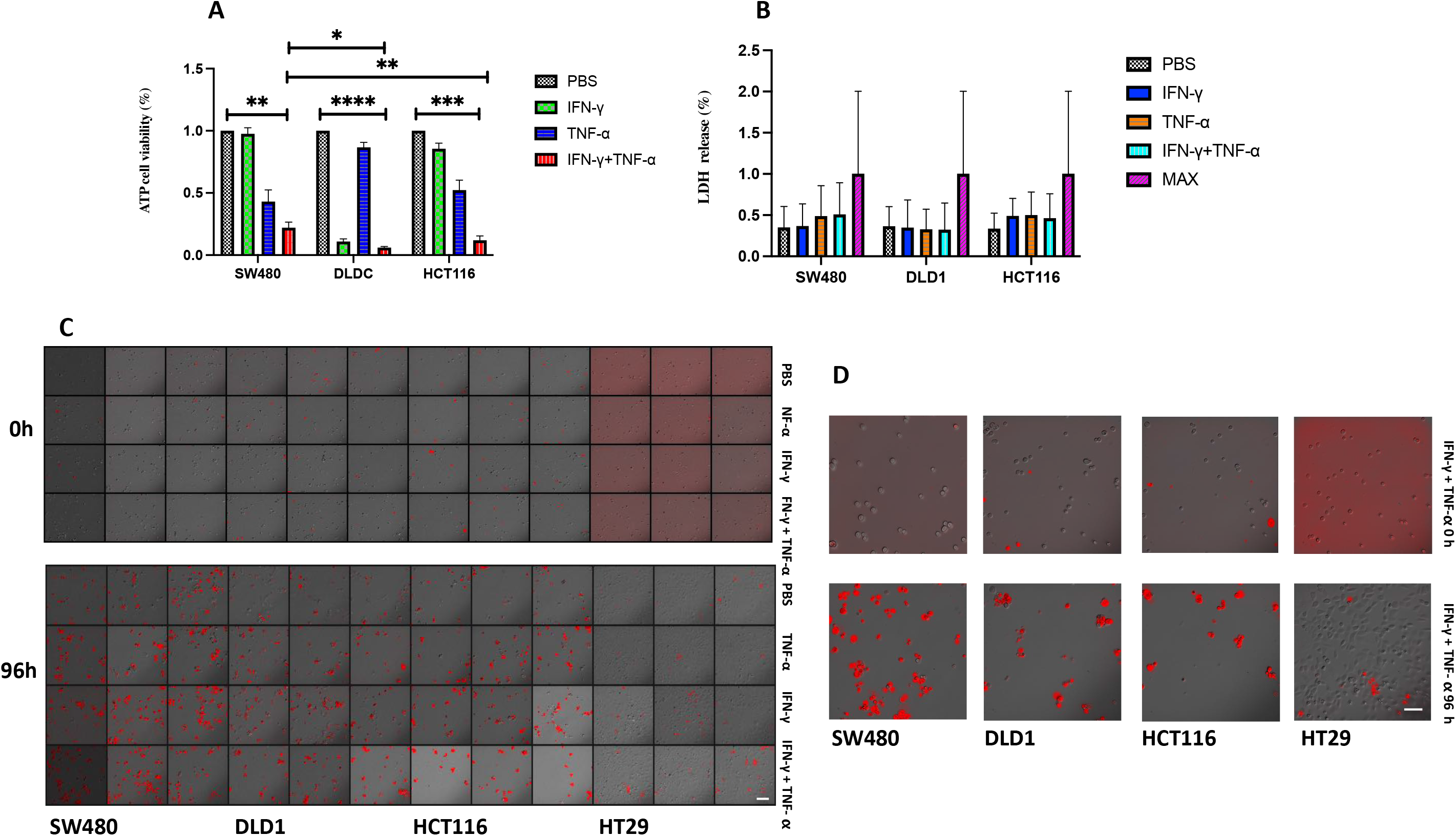
Treatment of IFN-*γ*, TNF-*α*, or IFN-*γ* + TNF-*α* induces PANoptosis. SW480, DLD1, HCT116 or HT29 were treated by PBS, IFN-γ, TNF-α, or IFN-γ + TNF-α for 96h. Cell death was measured by ATP-based cell viability (A), LDH release (B) and PI staining (C, D). Data are expressed as mean ± s.d. from three technical replicates. (mean ± s.e.m.). Two-tailed unpaired Student’s *t*-test was used to determine significance (\**P* < 0.05; \*\**P* < 0.01; \*\*\**P* < 0.001; \*\*\*\**P* < 0.0001; NS, not significant). Scale bars, 50 um.

We futher proceed the LDH release in SW480, DLD1 or HCT116 with IFN-γ, TNF-α or co-treatment of IFN-γ and TNF-α. TNF-α induced more LDH release in SW480 or HCT116 than IFN-γ, but co-treatment of IFN-γ and TNF-α induced most LDH release in SW480 or HCT116. And co-treatment of IFN-γ and TNF-α iuduced more LDH release in HCT116 than in SW480, the result was consistent with above ATP cell viability, co-treatment of IFN-γ and TNF-α induced PANoptosis including pyroptosis and apoptosis; but there was not different in inducing DLD1 death with PBS, IFN-γ, TNF-α or co-treatment of IFN-γ and TNF-α. (Figure 1B).

### Co-treatment of IFN-*γ* and TNF-*α* induces PANoptosis of SA-b-gal-positive (blue) cell

Co-treatment of IFN-γ and TNF-α can induce senescence in cells, we analysed whether senescent cells was sensitive to co-treatment of IFN-γ and TNF-α to induce cell death(18–20). After co-treatment of IFN-γ and TNF-α, SA-b-gal-positive (blue) cells reduced, it showed co-treatment of IFN-γ and TNF-α would induced PANoptosis of SA-b-gal-positive (blue) cell (Figure 2A), death rate of SA-b-gal-positive (blue) cells was positive relation with sensitivity to co-treatment of IFN-γ and TNF-α. The number of death cells in DLD1, HCT116, RKO was more than in SW480, SW620, and DLD1, HCT116, RKO is dMMR cancer cell lines, SW480, SW620 is pMMR cancer cell lines, it showed that dMMR cancer cell lines were more sensitive than pMMR cancer cell lines to co-treatment of IFN-γ and TNF-α. It suggested that the signal path in senescence is cross with the signal path in PANoptosis (Figure 2B, 2C).

**Figure 2.**
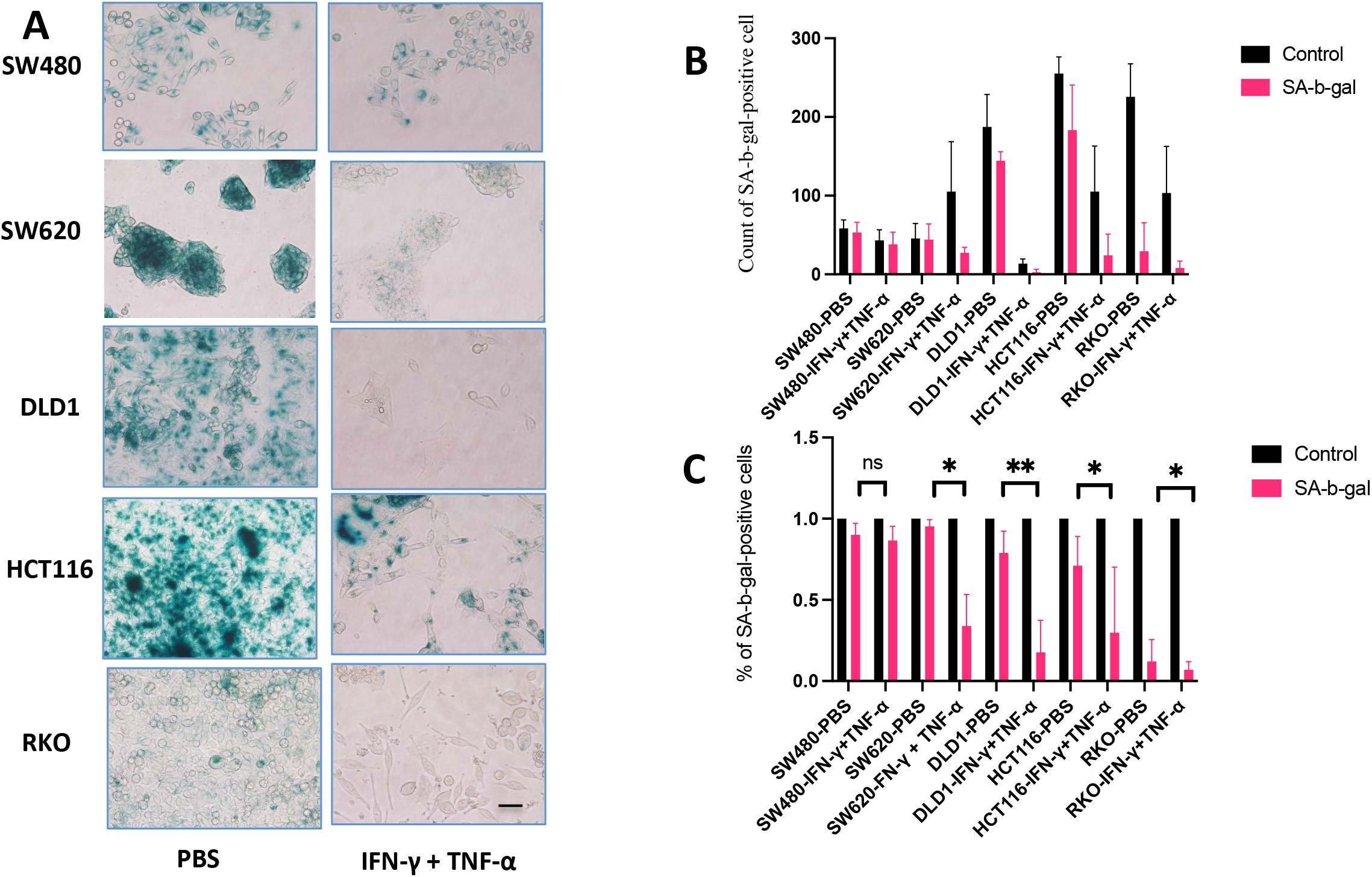
Co-treatment of IFN-*γ* and TNF-*α* induces PANoptosis of SA-b-gal-positive (blue) cell. The PANoptosis of SA-b-gal-positive (blue) cell in SW480, SW620, DLD1, HCT116 or RKO was induced by PBS or IFN-γ + TNF-α (A, B, C). SA-b-gal, % of SA-b-gal-positive (blue) cells. Data are expressed as mean ± s.d. from three technical replicates. (mean ± s.e.m.). Two-tailed unpaired Student’s *t*-test was used to determine significance (\**P* < 0.05; \*\**P* < 0.01; \*\*\**P* < 0.001; \*\*\*\**P* < 0.0001; NS, not significant). Scale bars, 50 um.

### Effect of of pan-caspase inhibitor on PANoptosis with co-treatment of IFN-*γ* and TNF-*α*

We futher analyzed effect of caspase inhibitors on PANoptosis induced by co-treatment of IFN-γ and TNF-α. The results showed that most cancer cell lines death was induced by co-treatment of IFN-γ and TNF-α, in which cell death was significantly induced in MC38 and HCT116, which were close to complete death after co-treatment of IFN-γ and TNF-α for 96 hours. The pan-caspase inhibitor Emricasan alone would not inhibit cell death, along with the same growth rate of cells comparing with the group treated with PBS, but it could promote cell proliferation in MC38; Emricasan suppressed cell death induced by co-treatment of IFN-γ and TNF-α, especially completely inhibiting cell death in MC38 or *Mlh1* KO CT26 co-treated with IFN-γ and TNF-α. Emricasan also inhibited cell death in HCT116 with co-treatment of IFN-γ and TNF-α, especially inhibiting 50% of cell death. During the experiment, we observed that emricasan also completely inhibited cell death in HCT116 after co-treatment of IFN-γ and TNF-α, but the cell activity was only 50% of that of control group with PBS, which was due to inducing dormancy in HCT116 after co-treatment of IFN-γ plus TNF-α plus Emricasan; Emricasan could not suppress cell death in SW620 with co-treatment of IFN-γ and TNF-α, which also explained that in addition to inducing caspase activation and cell death by co-treatment of IFN-γ and TNF-α, there were other factors to induce cell death such as inducing necrosis or activating p-H2A to induce genome damage(21) (Figure 3A, 3B). We further analyzed the effects of other inhibitors on cell death. After co-treatment of IFN-γ and TNF-α for 96 hours, a large number of cell death was induced, and the caspase-8 inhibitor Z-IETD-FMK could partially prevent cell death in MC38, *Mlh1* KO CT26, SW620 or HCT116, especially inhibiting 50% cell death in HCT116 and RKO, indicating that caspase-8 plays an important role on cell death in HCT116 and RKO induced by co-treatment of IFN-γ and TNF-α, but Z-IETD-FMK could not inhibit the cell death of vector control CT26 induced by co-treatment of IFN-γ and TNF-α; co-treatment of IFN-γ plus TNF-α plus Z-IETD-FMK plus Necrosulfonamide could superimpose on the inhibition of Z-IETD-FMK on cell death induced by co-treatment of IFN-γ and TNF-α in MC38, vector control CT26, *Mlh1* KO CT26, but not in SW620, HCT116 and RKO. MLKL inhibitor Necrosulfonamide could inhibit cell death induced by co-treatment of IFN-γ and TNF-α in MC38, vector control CT26, *Mlh1* KO CT26, SW620, HCT116 or RKO, among which the inhibition is the most significant in SW620, indicating that the necrosis pathway plays an important role on cell death of SW620 induced by co-treatment of IFN-γ and TNF-α (Figure 3C, 3D).

**Figure 3.**
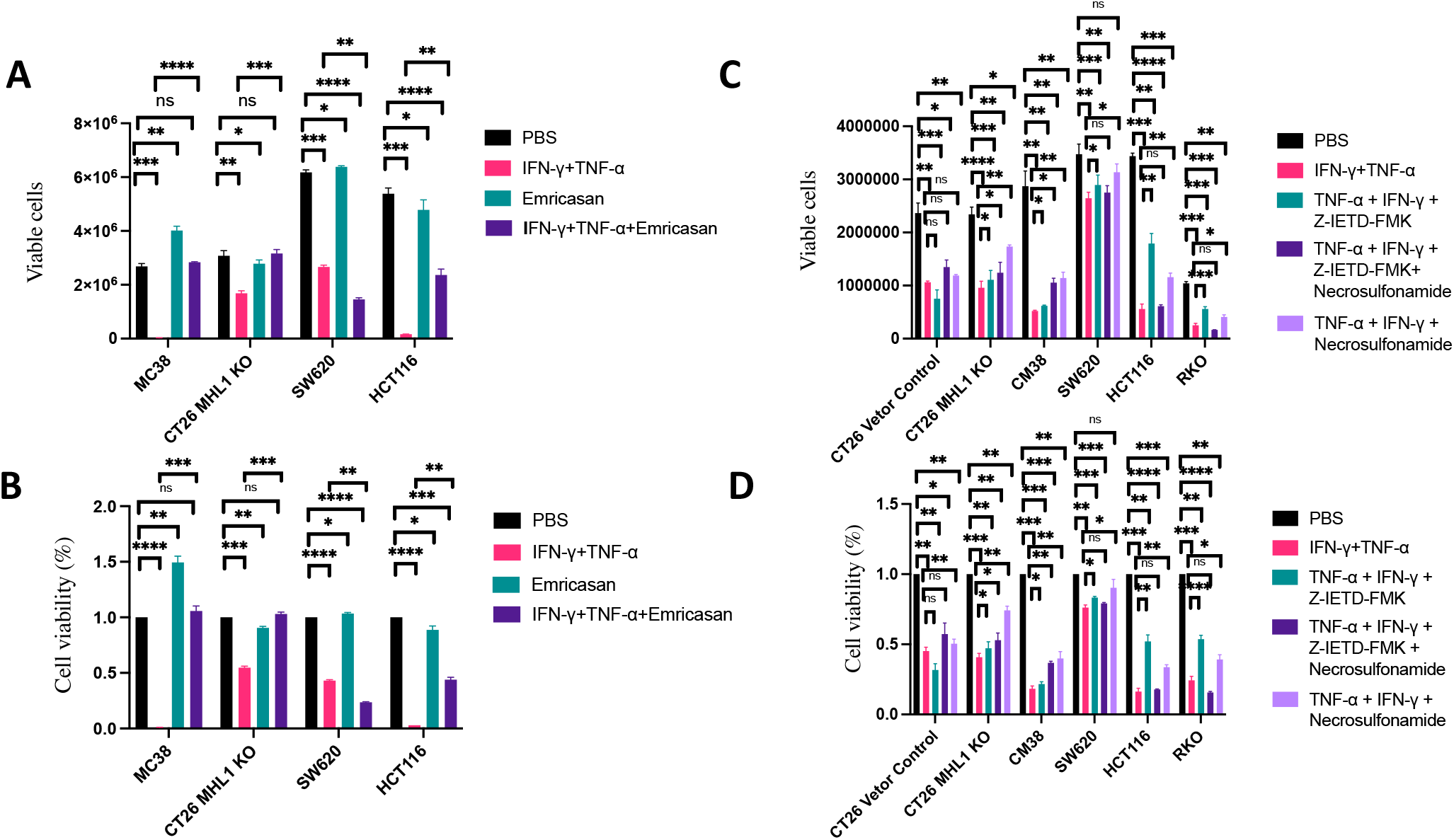
Co-treatment of TNF-*α*, IFN-*γ*, Emricasan (IDN-6556), Z-IETD-FMK and Necrosulfonamide induces PANoptosis. (A-B) MC38, CT26-6, SW620, HCT116 were treated with PBS, IFN-γ + TNF-α, Emricasan, or IFN-γ + TNF-α + Emricasan; (C-D) PBS, IFN-γ + TNF-α, IFN-γ + TNF-α + Z-IETD-FMK, IFN-γ + TNF-α + Z-IETD-FMK+Necrosulfonamide, IFN-γ + TNF-α + Necrosulfonamide to induce PANoptosis;. Viable cells were counted by Countstar Automated Cell Counter, % of Viable cells were shown. Data are expressed as mean ± s.d. from three technical replicates. (mean ± s.e.m.). Two-tailed unpaired Student’s *t*-test was used to determine significance (\**P* < 0.05; \*\**P* < 0.01; \*\*\**P* < 0.001; \*\*\*\**P* < 0.0001; NS, not significant).

### Caspase inhibitor Emricasan can inhibit the PANoptosis induced by co-treatment of IFN-*γ* and TNF-*α*

We next identified specific targeting of these molecules in PANoptosis induced by co-treatment with IFN-γ plus TNF-α plus Emricasan. The results showed that DLD1, HCT116 GSDMD could also be activated and cleaved by PBS treatment. The activated fragment P30 was naturally hyperactivated, but there was no activated fragment P30 in the cell membrane, so there was no polymer forming on the cell membrane, no forming pores, and no inducing cell death. 100 ng IFN-γ and 10 ng TNF-α increased the activation of GSDMD in the cytoplasm in DLD1 or HCT116 and produced more P30 fragments, emricasan with a final concentration of 20 μ M could not inhibit the activation of GSDMD in DLD1 with co-treatment of IFN-γ and TNF-α, but could in HCT116; while IFN-γ and TNF-α could also increase the activation of GSDMD on the cell membrane in DLD1 or HCT116, and emricasan could not inhibit the activation of GSDMD on the cell membrane in DLD1 with co-treatment of IFN-γ and TNF-α, but could do in HCT116, which was the main reason that DLD1 or HCT116 cells can be induced to pyrosis through activated P30 of GSDMD, forming pores in the cell membrane, inducing cell lysis; IFN-γ and TNF-α could also increase the activation of GSDMD in the nucleus in DLD1, but emricasan could not inhibit the activation of GSDMD in the nucleus in DLD1 with co-treatment of IFN-γ and TNF-α; however, emricasan could inhibit the activation of GSDMD in the nucleus in HCT116 with co-treatment of IFN-γ and TNF-α (Figure 4A, 4C).

**Figure 4.**
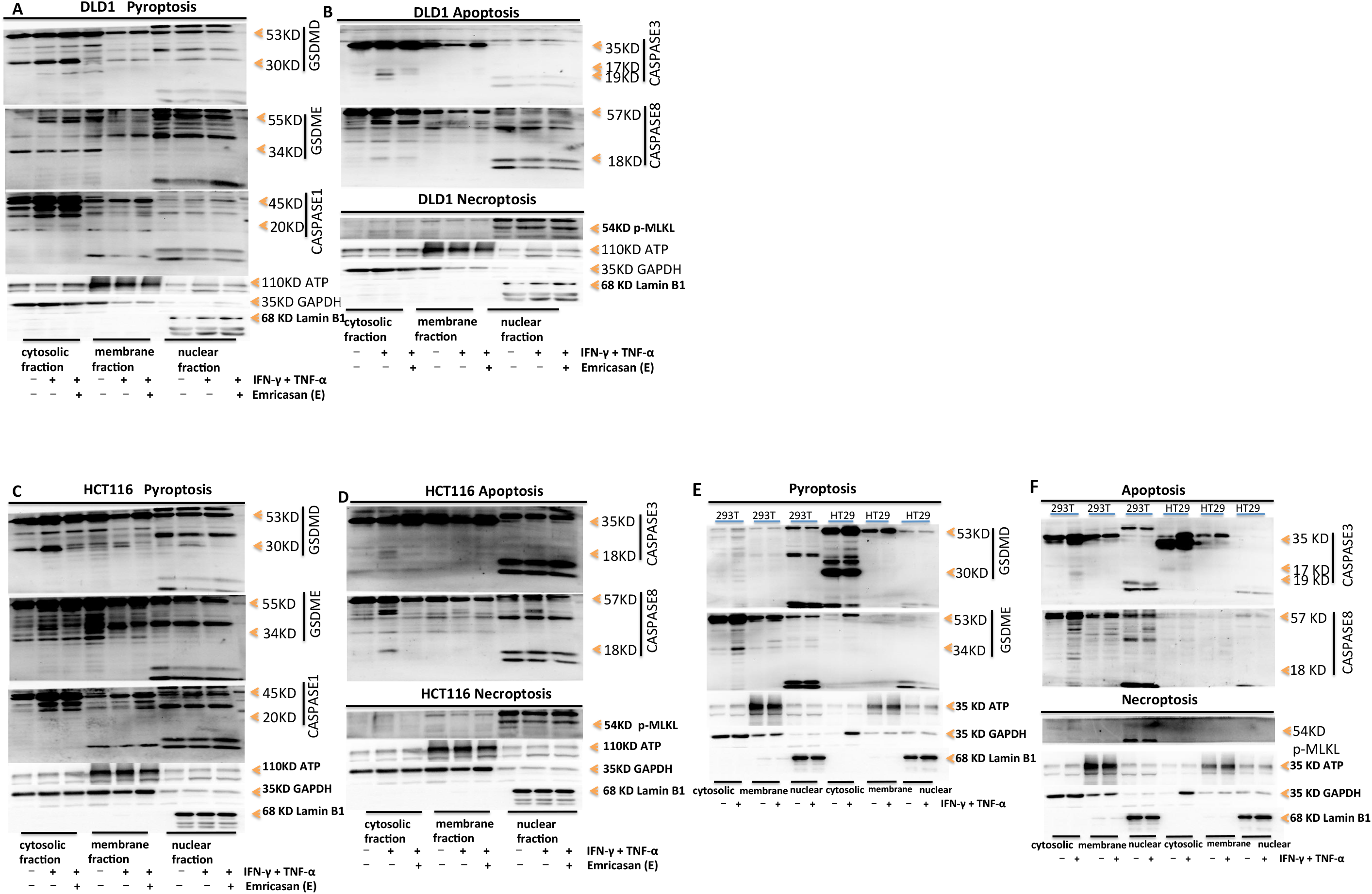
Co-treatment of IFN-*γ* + TNF-*α* + Emricasan (IDN-6556) induces PANoptosis. Immunoblot analysis of pro-(P53), activated (P30) GSDMD, pro-(P53) and activated (P34) GSDME, pro-(P45) and activated (P20) CASP1; pro-(P35) and cleaved (P19 and P17) CASP3, pro-(P55) and cleaved (P18) CASP8, phosphorylated MLKL (pMLKL) in DLD1 (A, B), HCT116 (C, D), 293T or HT29 (E, F) after co-treatment with IFN-γ + TNF-α or co-treatment with IFN-γ + TNF-α + Emricasan (IDN-6556) for 24h. ATP, GAPDH, Lamin B1 was used as membrane, cytosolic, nuclear internal control respectively. Data are representative of at least three independent experiments.

Since DLD1 does not express GSDME, which is consistent with other research reports, it was not analyzed here. HCT116 GSDME could also be activated and cleaved by the treatment of PBS. The activated fragment p34 was naturally hyperactivated. At the same time, the activated fragment p34 was also present in the cell membrane. Therefore, polymers were formed on the cell membrane to form pores, but the cells survived at the same time. This is called the natural hyperactivation of GSDME. IFN-γ plus TNF-α increased the activation of GSDME in the cytoplasm in HCT116, GSDME was cleaved into the P34 fragment, and emricasan with a final concentration of 20 μM inhibited the activation of GSDME by IFN-γ and TNF-α in the cytoplasm. IFN-γ and TNF-α could also induce the activation of GSDME in the cell membrane in HCT116, and emricasan also inhibited the activation of GSDME on the cell membrane in HCT116 with IFN-γ and TNF-α, which was the main reason that HCT116 cells could be induced to pyrosis through the activated GSDME p34, forming pores in the cell membrane, inducing cell lysis and causing cell pyrosis; IFN-γ and TNF-α could not induce the activation of GSDME in the nucleus in HCT116, so emricasan could not inhibit the activation of GSDME in the nucleus in HCT116 with IFN-γ and TNF-α (Figure 4C).

IFN-γ and TNF-α induced the activation of caspase-1 in the cytoplasm in DLD1 or HCT116, caspase-1 was cleaved into the activated P20 fragment. Caspase-1 was mainly expressed in the cytoplasm and a little in the cell membrane and nucleus. Emricasan could not inhibit the activation of caspase-1 in the cytoplasm in DLD1 with co-treatment of IFN-γ and TNF-α, indicating that emricasan could not inhibit pyroptotic cell death induced by co-treatment with IFN-γ and TNF-α; but Emricasan could inhibit the activation of caspase-1 in the cytoplasm in HCT116 with co-treatment of IFN-γ and TNF-α, indicating that emricasan could inhibit pyroptotic cell death in HCT116 induced by co-treatment of IFN-γ and TNF-α (Figure 4A, 4C). IFN-γ and TNF-α induced the activation of caspase-3 in the cytoplasm in DLD1 or HCT116, caspase-3 was cleaved into the active fragments, p17 and P19, and emricasan inhibited the activation of caspase-3 in the cytoplasm in DLD1 or HCT116 with co-treatment of IFN-γ and TNF-α; caspase-3 in DLD1 or HCT116 was mainly expressed in the cytoplasm and a little in the membrane. IFN-γ and TNF-α could not induce the activation of caspase-3 in the cell membrane in DLD1 or HCT116, and emricasan treatment could not induce the activation of caspase-3 in the membrane in DLD1 or HCT116 cell; there was no expression of caspase-3 in the nucleus in DLD1 or HCT116 and no activation of caspase-3. IFN-γ and TNF-α induced the activation of caspase-8 in the cytoplasm in DLD1 or HCT116, caspase-8 was cleaved into the active P18 fragment, and emricasan inhibited the activation of caspase-8 in the cytoplasm in DLD1 or HCT116 with co-treatment of IFN-γ and TNF-α; caspase-8 was expressed on the DLD1 or HCT116 cell membrane, Emricasan could not induce the activation of caspase-8 in the cell membrane in DLD1 or HCT116, so emricasan had no effect on activation of caspase-8 in the cell membrane in DLD1 or HCT116; caspase-8 was not expressed in the nucleus in DLD1 or HCT116 (Figure 4B, 4D). p-MLKL was induced in DLD1 or HCT116 with treatment of PBS, IFN-γ and TNF-α or Emricasan plus IFN-γ plus TNF-α, but p-MLKL was only activated in the nucleus, and there was no difference in activation level among the treatment of PBS, IFN-γ and TNF-α or Emricasan plus IFN-γ plus TNF-α. Emricasan could not inhibit p-MLKL activation in DLD1 or HCT116, indicating that p-MLKL was not activated through caspase pathway. In addition, there was no activated p-MLKL in the cytoplasm or membrane in DLD1 or HCT116, indicating that IFN-γ and TNF-α could not induce cell death by necrosis (Figure 4B, 4D). IFN-γ and TNF-α induced PANoptosis by pyrosis and apoptosis, but not by necrosis. Emricasan treatment could inhibit pyrosis and apoptosis, but had no effect on necrosis to induce PANoptosis.

We further identified specific molecules of PANoptosis in 293T or HT29 with co-treatment of IFN-γ and TNF-α. It showed that GSDMD is not expressed in 293T. GSDMD in HT29 could also be activated and cleaved by PBS treatment. The activated fragment P30 is naturally hyperactivated, but there was no activated fragment P30 in the cell membrane, so it did not form polymers on the cell membrane, forming pores, and inducing cell death. 100 ng IFN-γ and 10 ng TNF-α treatment induced the activation of GSDMD in the cytoplasm in HT29 and cleaved GSDMD into the P30 fragment, but IFN-γ and TNF-α could not induce the activation of GSDMD in the cell membrane and nucleus in HT29, so it could not form polymers on the cell membrane, forming pores, and inducing cell death. GSDME was not expressed in HT29. IFN-γ and TNF-α treatment could induce the activation of GSDME in the cytoplasm in 293T and cleave GSDME into the p34 fragment, but it could not induce the activation of GSDME in the cell membrane and nucleus. Therefore, it could not form polymers on the cell membrane, forming pores, and inducing cell death. IFN-γ and TNF-α treatment induced activation of caspase-3 in the cytoplasm in 293T, but could not induce activation of caspase-3 in the cell membrane and nucleus in 293T; IFN-γ and TNF-α could not induce the activation of caspase-3 in the cytoplasm, membrane and nucleus in HT29; IFN-γ and TNF-α treatment induced activation of caspase-8 in the cytoplasm in 293T, but a little level of activation; IFN-γ and TNF-α could not induce activation of caspase-8 in the cell membrane and nucleus in 293T; HT29 did not express caspase-8; thus, IFN-γ and TNF-α treatment did not induce PANoptosis by apoptosis in 293T or HT29. IFN-γ and TNF-α treatment will not induce the activation of p-MLKL in the cytoplasm, membrane and nucleus in 293T or HT29, so it could not induce PANoptosis by necrosis. IFN-γ and TNF-α treatment could not induce activation of GSDMD in the cell membrane in HT29 and activation of GSDME in the cell membrane in 293T, so it could not induce pyrosis to induce cell death, could not induce activation of caspase-3 and caspase-8 in 293T or HT29 to induce apoptosis, and could not induce activation of p-MLKL in 293T or HT29 to induce cell necrosis. Therefore, IFN-γ and TNF-α treatment could not induce PANoptosis in 293T or HT29 (Figure 4E, 4F).

### GSDMD, GSDME oligomerize into pore formation

To examine GSDMD oligomerization, which correlates with pore formation, we used a protocol that NT-GSDMD distinguishes monomeric and oligomeric GSDMD. Specifically, NT-GSDMD is oligomers under SDS-PAGE electrophoresis without a reducing agent, but oligomers become monomers after adding a reducing agent. The results showed that DLD1 and HCT116 GSDMD was activated by PBS treatment, and cleaved into P30 fragment. There was natural hyperactivation, and the P30 fragments existed in the cytoplasm, membrane and nucleus. IFN-γ and TNF-α enhanced the activation of GSDMD in the cytoplasm, membrane and nucleus in DLD1 or HCT116; under the non-reduced SDS-PAGE electrophoresis, the activated P30 fragment of GSDMD was a polymer with a molecular weight of more than 400kd in DLD1 or HCT116. The pore of P30 polymers that existed in the cytoplasm and membrane, but not in the nucleus in DLD1, was present in the concentration gel and the separation gel. The pore of P30 polymers existed in the cytoplasm, membrane and nucleus in HCT116. Noted that GSDMD was activated and cleaved into P30, and the pores of 16 P30 polymers were formed on the cell membrane to induce cell death in DLD1 or HCT116 with treatment of PBS or IFN-γ and TNF-α. DLD1 did not express GSDME. After the treatment with PBS or IFN-γ and TNF-α on HCT116, GSDME was activated and cleaved into p34 fragment in HCT116. The p34 fragment existed in the cytoplasm, membrane and nucleus, and IFN-γ and TNF-α enhanced activation of GSDME in HCT116; under the non-reduced SDS-PAGE electrophoresis, activated p34 fragment of HCT116 GSDME formed a polymer with a molecular weight of more than 400kd. As the pores of 16 P34 polymers that existed in the cytoplasm and membrane, but not in the nucleus in HCT116, were present in the concentrated gel and the separation gel, indicating that HCT116 could form polymer pores in the cell membrane and induce cell death (Figure 5A). p-MLKL in DLD1 or HCT116 with treatment of PBS or IFN-γ and TNF-α could be activated, and the level of activation was not different, but only existed in the nucleus, and formed polymers only in the nucleus, but not on the cell membrane to induce cell necrosis (Figure 5B). This was also consistent with the previous results that IFN-γ and TNF-α treatment could induce the activation of GSDMD and GSDME in DLD1 or HCT116, inducing the formation of multimeric pores on the cell membrane to promote PANoptosis through pyroptosis and apoptosis, but not necrosis.

**Figure 5.**
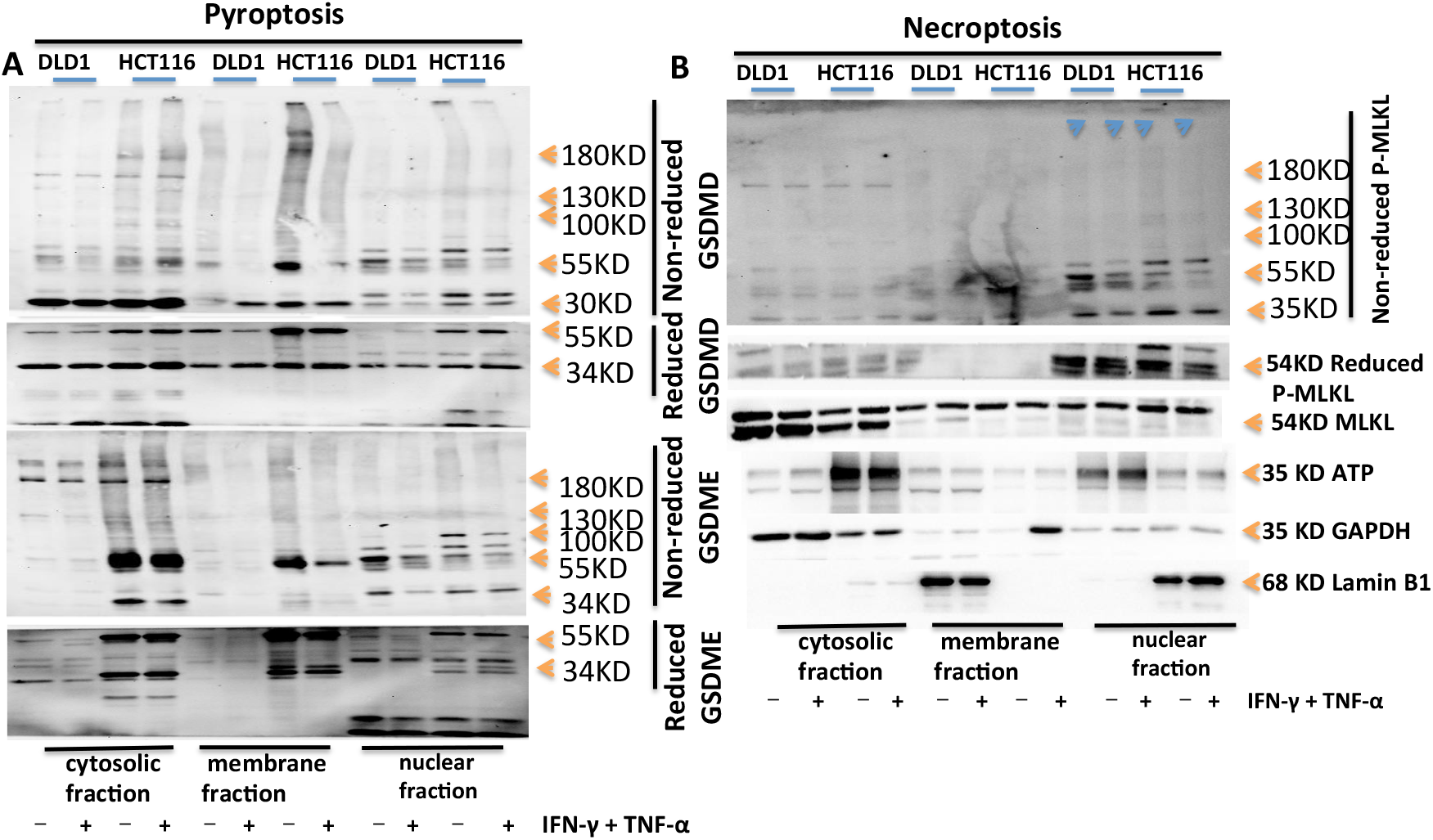
Oligomerization of NT-GSDMD, NT-GSDME and p-MLKL triggers PANoptosis after co-treatment of IFN-*γ* + TNF-*α*. (A–B) Immunoblot analysis of (A) pro-(P53), activated (P30), oligomerized GSDMD, and pro-(P53), activated (P34), oligomerized GSDME, (B) tMLKL, phosphorylated MLKL (pMLKL) and oligomerized pMLKL induced by PBS or IFN-γ + TNF-α in DLD1 or HCT116; the activative NT-GSDMD, NT-GSDME, p-MLKL could formed oligomerer, 100kD, 130KD, 180KD, and bigger oligomerer. The activative NT-GSDMD, NT-GSDME mainly in cytosolic fraction and membrane fraction, littlely in nuclear fraction; the activative p-MLKL mainly in nuclear fraction, littlely in cytosolic fraction and membrane fraction. ATP, GAPDH, Lamin B1 was used as membrane, cytosolic, nuclear internal control respectively. Data are representative of at least three independent experiments.

### DNA sensor for DNA fragments in Cancer cells with dMMR is in nucleus, not in cytosolic

When MHL1 was absent, cancer cells’ genome was unstable, releasing many DNA fragments and promoting many genes to express. We analysed expression and location of DNA sensors in cancer cells. It showed that cGAS, AIM2, ZBP1 or ASC in DLD1 or HCT116 with treatment of PBS or IFN-γ and TNF-α, had no difference in the expression level, but cGAS was only expressed in DLD1 but not in HCT116; after the treatment of IFN-γ and TNF-α in HCT116, lamin B1 was activated and cleaved into P45 fragment, indicating that IFN-γ and TNF-α activated apoptotic molecules caspase-3 and caspase-7 to degrade the nuclear fiber layer and induce apoptosis; However, after the treatment of PBS in HCT116, it also showed lamin B1 activation and to be cleaved into P45 fragment, indicating that HCT116 had natural nuclear membrane damage, which was consistent with the natural activation of caspase-7 (Figure 6A); however, in 293T, both the treatment of PBS and IFN-γ plus TNF-α could induce ZBP1 to be expressed, and the level of expression was not different, all of them were only in the nucleus, but ZBP1 was not expressed in HT29, which was consistent with the result that p-MLKL was not naturally activated in HT29; both the treatment of PBS and IFN-γ plus TNF-α could induce cGAS to be expressed in 293T or HT29 but cGAS only existed in the nucleus in HT29, and cGAS was expressed in the cytoplasm and nucleus in 293T; AIM2 is not expressed in HT29, which indicates that the previous natural hyperactivation of GSDMD cannot be activated by AIM2 pathway, but by other pathways; both the treatment of PBS and IFN-γ plus TNF-α could induce AIM2 to be expressed in 293T, AIM2 only existed in the nucleus; in addition, lamin B1 was not activated in 293T or HT29 and was not cleaved into P45 fragment, which was similar to the previous result that IFN-γ and TNF-α did not induce apoptosis in 293T or HT29 (Figure 6B).

**Figure 6.**
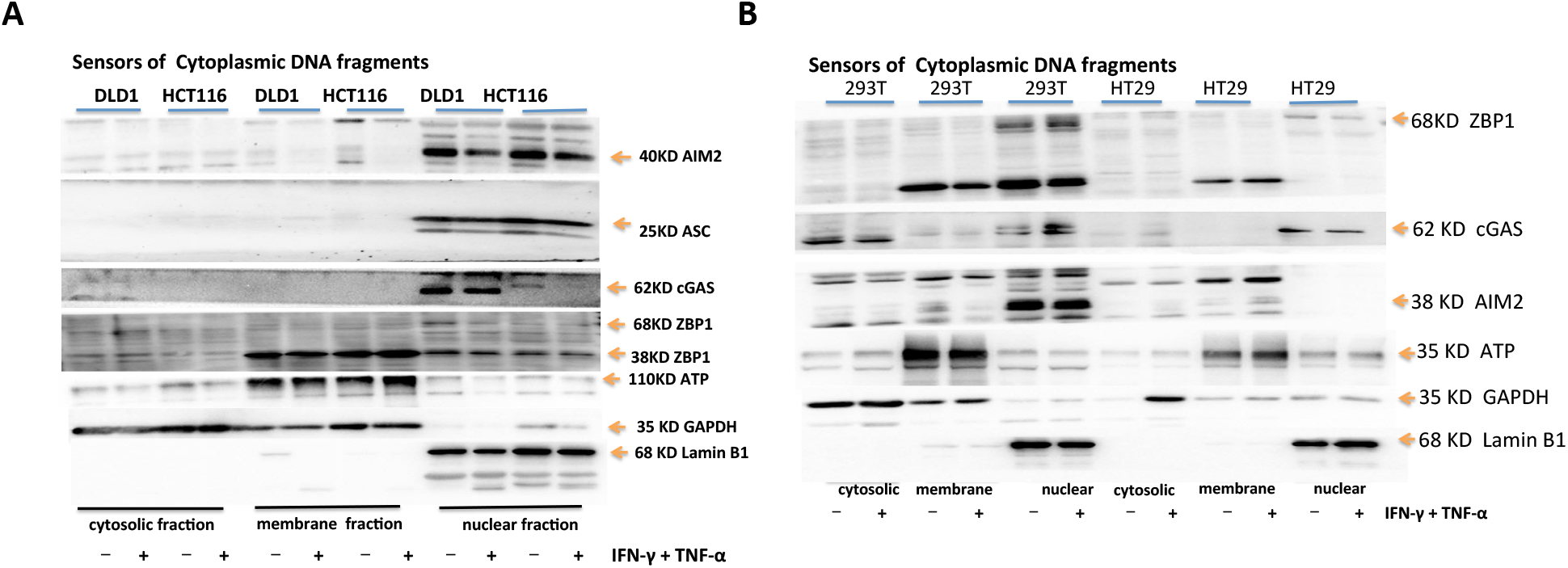
DNA sensor for DNA fragments in Cancer cells with dMMR is in nucleus, not in cytosolic. (A–B) Immunoblot analysis of cGAS, ZBP1, AIM2, ASC in DLD1, HCT116, 293T or HT29. ATP, GAPDH, Lamin B1 was used as membrane, cytosolic, nuclear internal control respectively. Data are representative of at least three independent experiments.

### Hyperactivation of PANoptosis effective moleculars by AIM2-ZBP-RIPK1-RIPK3-ASC-CASP8-CASP1 signal pathway

We further investigated effect of AIM2 or ZBP1 on hyperactivation of PANoptosis effective moleculars by RNA Interference. After treating DLD1 and HCT116 with siAIM2-1 and siAIM2-3 respectively, the activation of GSDMD p30, GSDME p34 and p-MLKL in DLD1 or HCT116 was inhibited, indicating that AIM2 was a receptor for nuclear DNA fragments in DLD1 or HCT116 cells. DLD1 and HCT116 genomes with mismatch repair deficiency were unstable, forming more nuclear DNA fragments. GSDMD p30, GSDME p34, p-MLKL were activated through AIM2-RIPK1-RIPK3-ASC-CASP8-CASP1, GSDMD p30, GSDME p34 were shown in Figure 4 that they were located in the cytoplasm and membrane, while p-MLKL was located in the nucleus. After processing DLD1 and HCT116 with si-ZBP1-3 and si-ZBP1-1 respectively, the result showed that si-ZBP1-3 and si-ZBP1-1 knocking down-the expression of ZBP1 in DLD1 and HCT116 respectively inhibited the activation of GSDMD p30 and p-MLKL in DLD1 and HCT116, indicating that ZBP1 was a receptor for nuclear DNA fragments in DLD1 and HCT116 cells. The DLD1 and HCT116 genomes with mismatch repair deficiency were unstable, forming more nuclear DNA fragments in cells. GSDMD p30 and p-MLKL were activated through ZBP1-RIPK1-RIPK3-ASC-CASP8-CASP1. GSDMD p30 was located in the cytoplasm and membrane, as shown in Figure 4, while p-MLKL was located in the nucleus (Figure 7A).

**Figure 7.**
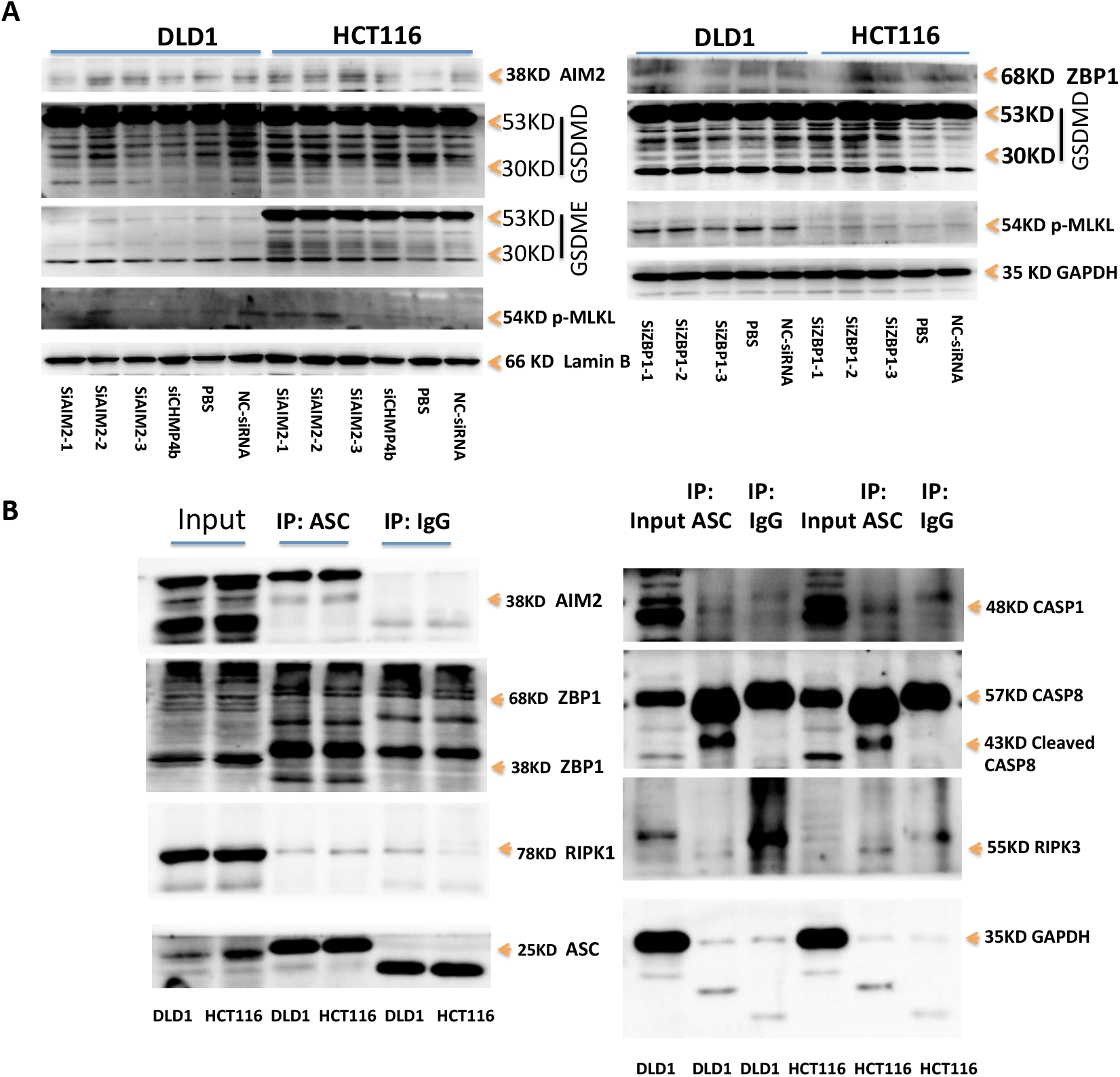
Hyperactivation of PANoptosis effective moleculars by AIM2-ZBP1-RIPK1-RIPK3-ASC-CASP8-CASP1 signal pathway. (A–D) Immunoblot analysis of AIM2, ZBP1, GSDMD, GSDME, p-MLKL in DLDL1 or HCT116 treated with siAIM2s or siZBP1s (A). Immunoblot analysis of AIM2, ZBP1, RIPK1, ASC, CASP1, CASP8, RIPK3 in DLDL1 or HCT116 immunoprecipitated with IgG control antibodies or anti-ASC antibodies (B). GAPDH was used as the internal control. Data are representative of at least three independent experiments.

### AIM2 and ZBP1 both act as DNA receptors and upstream regulators, which are required to form PANoptosome

We sought to understand the molecular relationship between AIM2 and ZBP1 in inducing inflammatory cell death and PANoptosis in DLD1 or HCT116. We observed interactions of ASC with AIM2, ZBP1, CASP1, CASP8, RIPK3, RIPK1 in DLD1 or HCT116 by immunoprecipitation. AIM2-ZBP1-ASC-RIPK1-RIPK3-CASP1-CASP8 could form a PANoptotic complex, activated CASP1 could cleave and activate GSDMD, activated CASP8 could activate CASP3, activated CASP3 could cleave and activate GSDME, and activated p-RIPK3 could directly activate p-MLKL, thereby activating GSDMD p30, GSDME p34, p-MLKL through AIM2-ZBP1-ASC-RIPK1-RIPK3-CASP1-CASP8 PANoptotic complex. The results showed that any knockdown of AIM2 or ZBP1 affected the activation of GSDMD p30, GSDME p34, and p-MLKL, indicating that AIM2 and ZBP1 both acted as DNA receptors and upstream regulators of the PANoptotic complex. The activated GSDMD p30 and GSDME p34 were shown in Figure 4, located in the cytoplasm and membrane, while p-MLKL was located in the nucleus (Figure 7B).

### The aggregation of N-GSDMD or N-GSDME is in the nucleus

Next, we sought to find out how pyroptotic molecules induced cell death after co-treatment of IFN-γ and TNF-α by time-lapse confocal images. The results showed that GSDMD-EGFP or GSDME-EGFP mainly existed in the cytoplasm and cell membrane in DLD1 or HCT116, and after the treatment of IFN-γ and TNF-α, most of them formed bright spots in the nucleus and formed aggregates, which was consistent with the previous result that the aggregation of N-GSDMD or N-GSDME was in the nucleus after the treatment of IFN-γ and TNF-α (Figure 8A, 8B, 8C). In addition, GSDMD-EGFP-expressing-DLD1 or GSDMD-EGFP or GSDME-EGFP expressing-HCT116 mostly had cell membrane and nucleus rupture. One of the reasons was that GSDMD-EGFP and GSDME-EGFP were induced to aggregate in the nucleus after cell death (Figure 8D, 8E).

**Figure 8.**
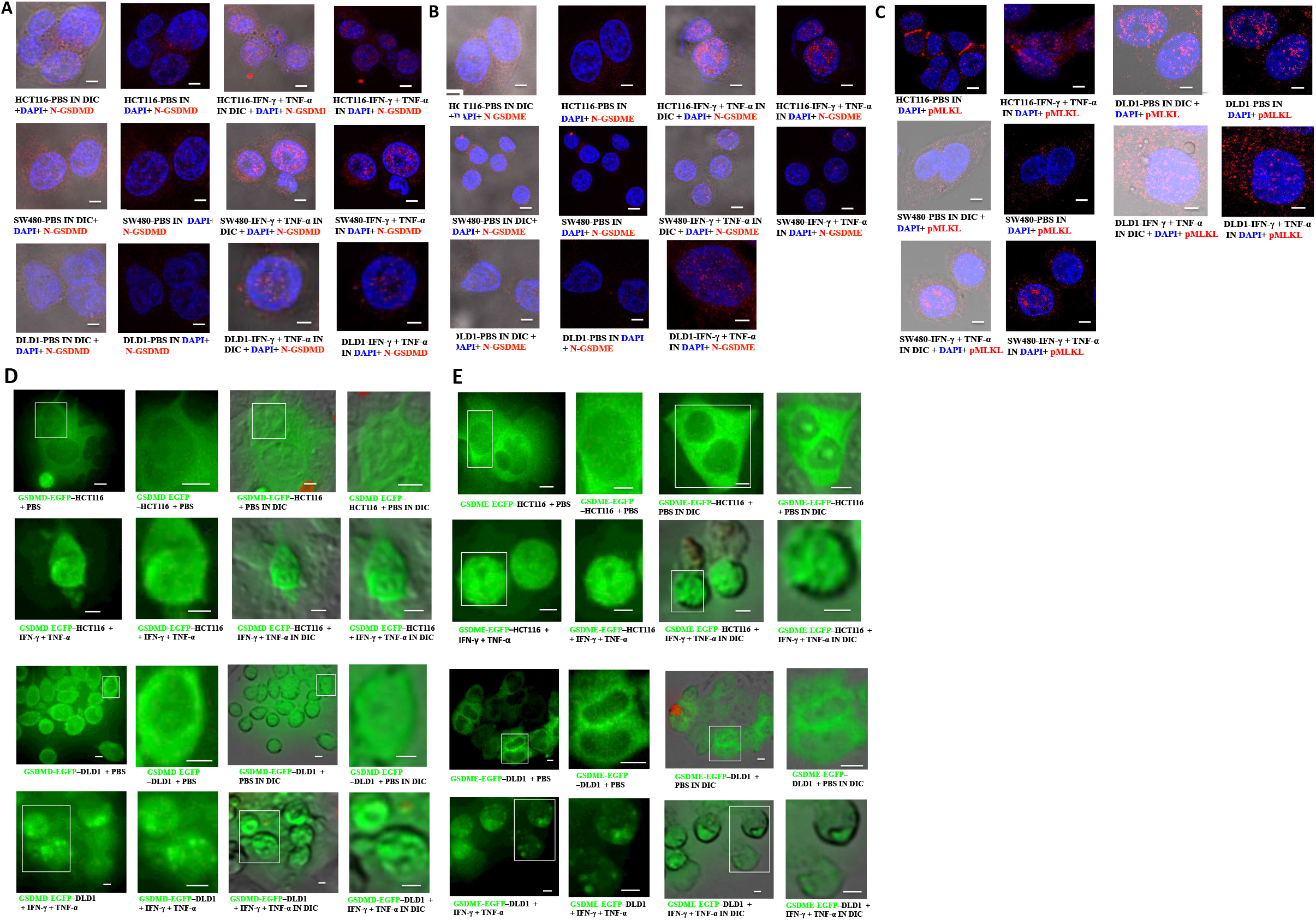
Location of PANoptotic molecules induced by co-treatment of IFN-*γ* + TNF-*α* using immunofluorescence and time-lapsed confocal microscopy. Fixed-cell microscopy images of NT-GSDMD(red), NT-GSDME(red), p-MLKL (red) in HCT116, SW480, DLD1 treated with PBS or IFN-γ + TNF-α (A, B, C) and time-lapse confocal images of either GSDMD-EGFP–expressing or GSDME-EGFP–expressing DLD1 or HCT116 cells after a 24-hours PBS or TNF-α + IFN-γ treatment (D and E). Nucleus stained by DAPI (bule), DIC, differential interference contrast. Results are representative of at least three independent experiments. Scale bars, 5 mm.

### Triton X-100 treatment on SW620, DLD1, HCT116 or HT29 induces cell outer membrane damage

High concentration of Triton X-100 can lyse cells, and low concentration of Triton X-100 can increase cell membrane permeability. Triton X-100 is used to simulate the effect molecules of T cell killing tumor cells, such as perforin and granzyme. When treatment with Triton X-100 at concentration of 0.01, 0.008, 0.006, 0.004 or 0.002% v / V in SW620, DLD1, HCT116 or HT29, it could immediately induce a large number of cell death. However, when treatment with Triton X-100 at the concentration of 0.001 % v / V, it could immediately induce a large number of cell death in HT29, but it could induce little cell death in SW620, DLD1 or HCT116. When treatment with Triton X-100 at the concentration of 0.001 % v / V for 7 hours, it could induce a large number of cell death in SW620 or HT29, but it could induce little cell death in DLD1 or HCT116, and SW620 and HT29 are mismatch repair proficient cell lines, but DLD1 and HCT116 are mismatch repair deficient cell lines. The reason was that DLD1 and HCT116 (dMMR) led to the formation of a large number of DNA fragments, activating AIM2-ZBP1-CASP1-CASP8-GSDMD-GSDME pathway to form pores on the cell membrane and causing cell membrane damage, finally inducing membrane repair system. Therefore, DLD1 or HCT116 (dMMR) was not sensitive to the low concentration of membrane damage agent Triton X-100, while SW620 or HT29 (pMMR) did not induce membrane repair system so that they were sensitive to cell membrane damage at the low concentration of Triton X-100(22) (Figure 9A, 9B, 9C, 9D).

**Fig 9.**
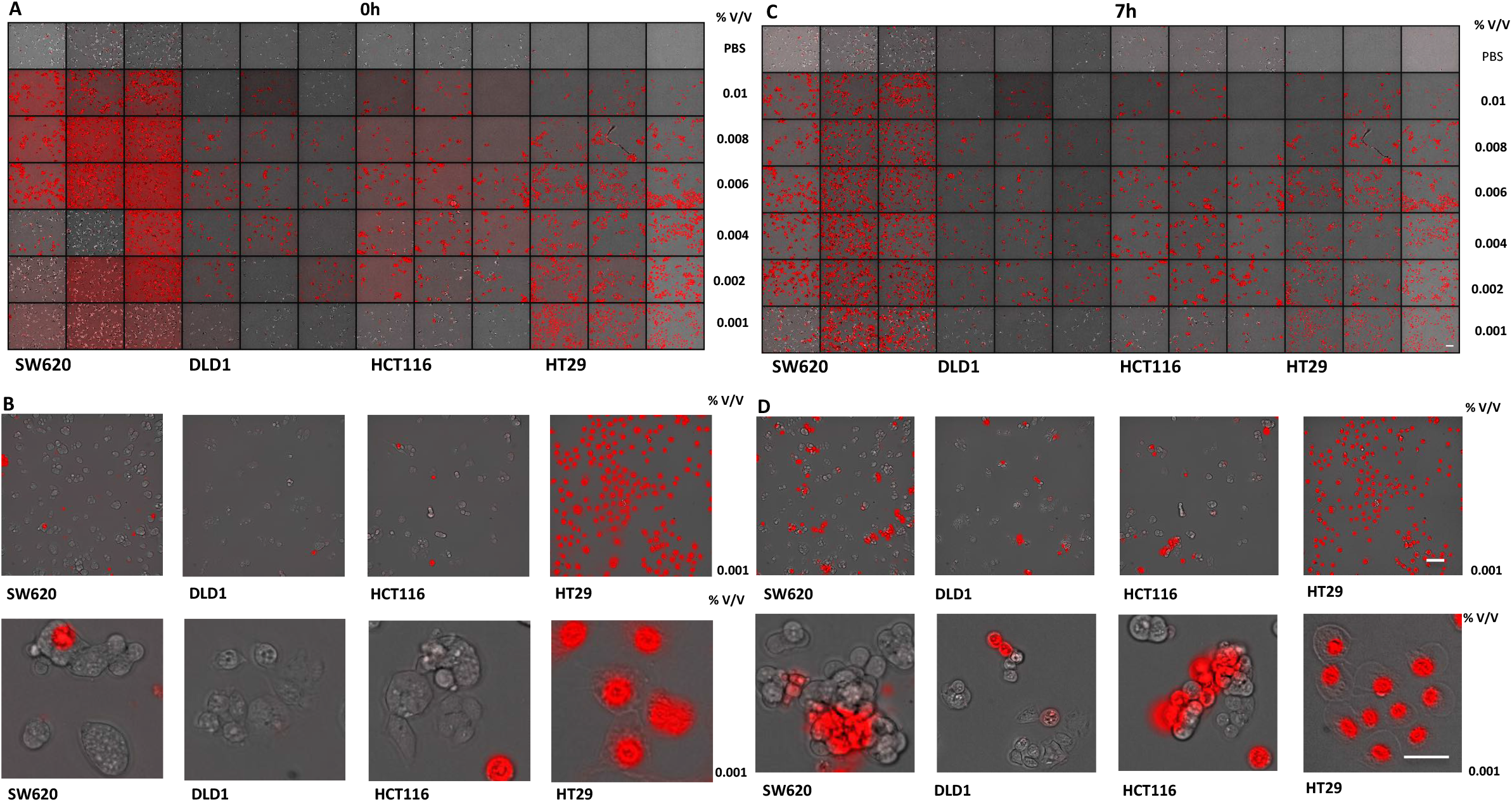
Triton X-100 induces lysis of cell membrance. SW620, DLD1, HCT116 or HT29 were treated by Triton X-100 at 0.01, 0.008, 0.006, 0.004, 0.002, 0.001 % v/v for 0 h, 7h. All cells was stained by PI, and time-lapse confocal images of PI +DIC. Scale bars, 50 um.

### *Mlh1* knockout CT26 undergoes natural hyperactivation of GSDMD, GSDME and p-MLKL

We further investigated the hyperactivation of of PANoptotic molecules in dMMR CT26 compared with pMMR CT26. Comparing GSDMD in WT CT26 with treatment of PBS, GSDMD in CT26 *Mlh1* KO1 or CT26 *Mlh1* KO2 with treatment of PBS was natural hyperactivation and cleavage of P30 fragments, and treatment with IFN-γ and TNF-α enhanced the activation of GSDMD and induced pyroptotic cell death; GSDMD in CT26 VT1 was natural hyperactivation in the presence of PBS; GSDME in CT26 *Mlh1* KO1 or CT26 *Mlh1* KO2 was also natural hyperactivation and cleavage of P34 fragments, comparing that in WT CT26 with treatment of PBS. There was no significant difference in activation of GSDME among treatment of PBS, TNF-α, IFN-γ or IFN-γ and TNF-α; GSDME in CT26 VT1 or CT26 VT2 was also natural hyperactivation in the presence of PBS; p-MLKL in CT26 *Mlh1* KO1 or CT26 *Mlh1* KO2 also showed significant natural hyperactivation with treatment of PBS, compared with that in WT CT26 with treatment of PBS, and the activation level was no significant difference among the treatment of PBS, TNF-α, IFN-γ or TNF-α and IFN-γ; At the same time, p-MLKL in CT26 VT1 or CT26 VT2 with treatment of PBS had also weak natural hyperactivation, the activation level was no significant difference among the treatment of PBS, TNF-α, IFN-γ or TNF-α and IFN-γ. It indicated that CT26 *Mlh1* KO led to the formation of more DNA fragments, thus activating the cytoplasmic receptors, such as cGAS, AIM2 and ZBP1. However, our previous results showed that these receptors existed in the nucleus, in which the nuclear dsDNA-AIM2-ZBP1-RIPK3-p-MLKL pathway activated p-MLKL and locked it in the nucleus, losing activation from the external signal stimulation, and leading to escape from cell necrosis. At the same time, there was also natural hyperactivation of dsDNA-AIM2-ZBP1-CASP1-CASP8-GSDMD-GSDME. At the same time, we found that CT26 VT1 and CT26 VT2 also exhibited natural hyperactivation of GSDMD, GSDME and p-MLKL, because they were constructed by cas9-expressing lentiviral vector, they could induce natural hyperactivation of GSDMD, GSDME and p-MLKL through viral infection. This was also the same conclusion of other studies that influenza virus infection can naturally activate ZBP1(23). However, our experimental results showed that the natural hyperactivation of p-MLKL in CT26 *Mlh1* KO was stronger than that in CT26 VT. The relevant conclusions can be further verified by the experiment of non viral vector *Mlh1* KO cell (Figure 10A, 10B).

**Fig 10.**
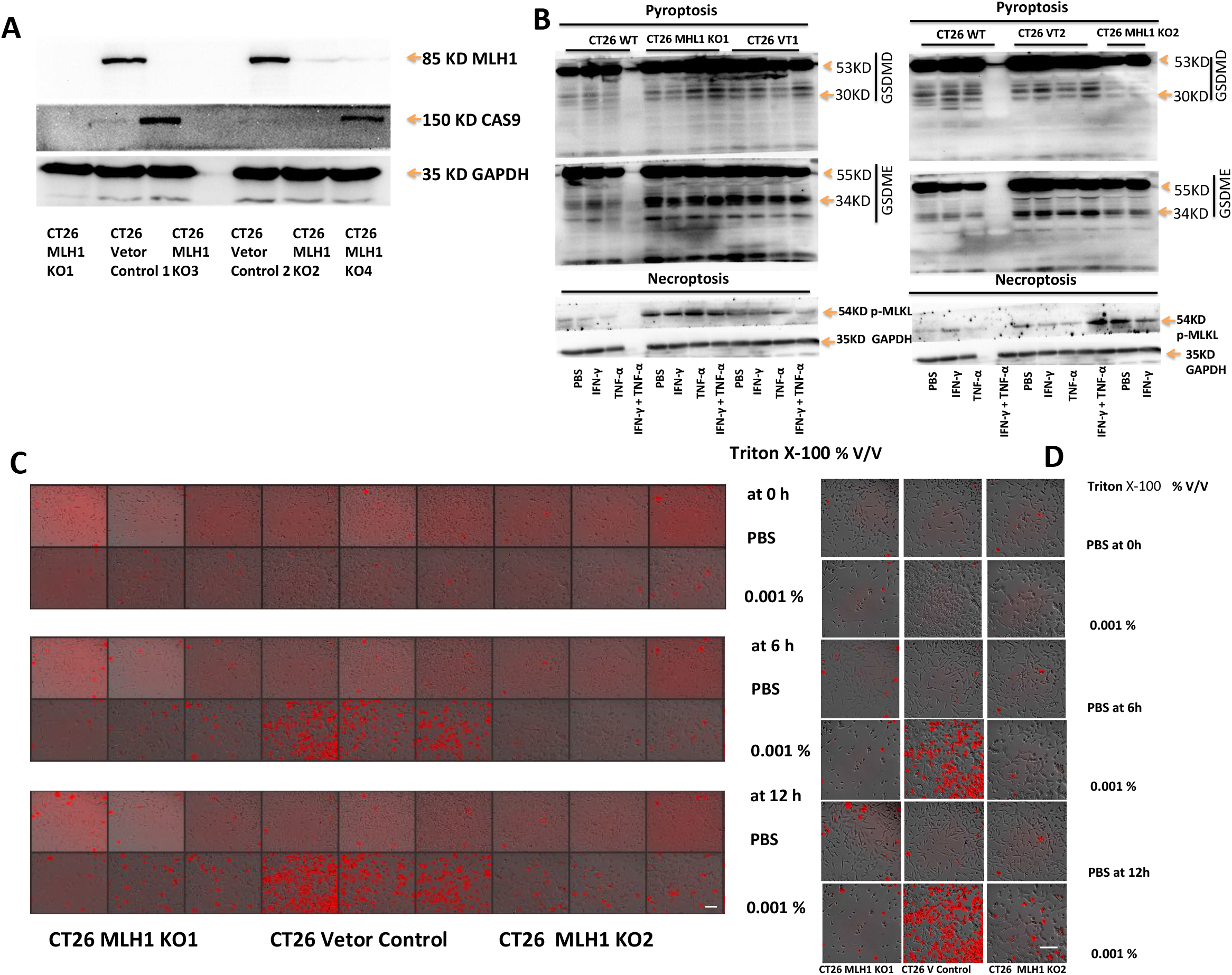
Triton X-100 induces lysis of cell membrance in CT26 MLH1 KO or CT26 Vetor Control. Immunoblot analysis of MHL1, CAS9 in CT26 Vetor Control, CT26 MHL1 KO (A); pro-(P53), activated (P30) GSDMD, pro-(P53), activated (P34) GSDME and p-MLKL in CT26 WT, CT26 MHL1 KO, CT26 Vetor Control, after treatment with IFN-γ alone, TNF-α alone, or co-treatment with IFN-γ + TNF-α for 96h(B). GAPDH was used as the internal. MLH1 KO1, CT26 Vetor Control, CT26 MLH1 KO2 were treated by Triton X-100 at 0.001 % v/v for 0 h, 6h, or 12h, all cells was stained by PI, and time-lapse confocal images of PI +DIC. Scale bars, 50 um.

### Triton X-100 treatment on *Mlh1* KO CT26 induces cell outer membrane damage

As shown above that *Mlh1* KO CT26 can repeat the natural hyperactivation of GSDMD, GSDME or p-MLKL in DLD1, HCT116 (dMMR), SW620 or HT29 (pMMR), so it can also repeat the sensitivity to Triton X-100. Triton X-100 at the concentration of 0.001 % v / V was used to treat CT26 *Mlh1* KO1, CT26 Vector Control or CT26 *Mlh1* KO2 for 6 hours or 12 hours, it could induce a large number of cell death in CT26 Vector Control, while it only induced little cell death in CT26 *Mlh1* KO1 or CT26 *Mlh1* KO2. This showed that CT26 cells with *Mlh1* gene knockout caused a large number of DNA fragments, which caused cell membrane damage through the AIM2-ZBP1-CASP1-CASP8-GSDMD-GSDME pathway, inducing the membrane repair mechanism, so that they were insensitive to the membrane damage agent Triton X-100. However, p-MLKL was mainly activated in the nucleus in CT26 *Mlh1* KO, so it could not damage the cell membrane and induce cell necrosis. In conclusion, the membrane damage induced by *Mlh1* knockout activated the membrane repair system through pyroptotic molecules GSDMD and GSDME, rather than necrotic molecule p-MLKL (Figure 10C, 10D).

## Discussion

Our experimental results showed that treatment alone with the concentration of TNF-α over 1ng could induce tumor cell death, and treatment alone with the concentration of IFN-γ up to 100 ng could not induce tumor cell death, but treatment alone with the concentration of IFN-γ at 100 ng could induce a large number of cell death in DLD1 and treatment alone with 10ng TNF-α just induced a little cell death, the mortality induced by the two treatments was very significant difference in DLD1, which was consistent with other studies. When caspase-8, RIPK1 or FADD is deficient, IFN-γ can induce cell death by ZBP1 signal pathway(24); only IFN-γ and TNF-α the highest cell death rate was induced by comparing that induced by IFN-γ alone or TNF-α alone, and the difference was very significant, and this result could be repeated in many cell lines, which indicates that the co-treatment of IFN-γ and TNF-α can induce PANoptosis including pyroptosis, apoptosis and necrosis, which can lead to higher cell death rate. At the same time, our result showed that the cell death rate induced by co-treatment of IFN-γ and TNF-α in DLD1, HCT116 and RKO cells with mismatch repair deficiency was higher than that in SW480, SW620 and HT29 cells with mismatch repair proficiency, and the difference was significant, which inspired us to study the mechanism. In DLD1 or HCT116, we could clearly see IFN-γ and TNF-α-induced significant activation of PANoptotic molecules, including pyroptotic molecules GSDMD, GSDME, apoptotic molecules caspase-3, caspase-7, caspase-8 and necrotic molecule p-MLKL, while GSDMD, GSDME was activated in the cytoplasm, caspase-3, caspase-7, caspase-8 was mainly in the cytoplasm, while p-MLKL was mainly in the nucleus, and there was no difference in the activation level of p-MLKL induced by between treatment of PBS and IFN-γ and TNF-α, indicating that p-MLKL is locked in the nucleus, and cell death cannot be induced by necrosis, but can only be induced by pyroptosis and apoptosis. Compared that with HT29 (mismatch repair proficiency), we found that HT29 had multiple defects in cell death pathways, GSDME was not expressed, caspase-8 was not expressed, and AIM2 was not expressed, which also explains that HT29 has no effect on treatment of IFN-γ and TNF-α, and is insensitive to cell death induced by treatment of IFN-γ and TNF-α, treatment of IFN-γ and TNF-α could induce activation of HT29 GSDMD and cleaved it into p30 fragment, but active p30 fragment only existed in the cytoplasm but not in the membrane, so that it cannot induce pyroptotic cell death and it can only induce weak activation of caspase-3 and caspase-8, so it can not induce apoptotic cell death, nor can it activate p-MLKL to induce necrotic cell death, so it can not induce cell death by PANoptosis.

At the same time, we observed treatment with IFN-γ and TNF-α in SW480, DLD1 or HCT116 induced N-GSDMD, N-GSMDE and p-MKLL to gather in the nucleus by immunofluorescence, forming bright patches and polymers. This is inconsistent with the polymerization of N-GSDMD and N-GSMDE on the cell membrane in other studies, which needs further study (25, 26).

In addition, DLD1 or HCT116 cells with mismatch repair dificiency released exonuclease 1, leading to increased single strand DNA formation, depletion of RPA, DNA breakage and abnormal DNA repair intermediates, eventually leading to chromosome abnormalities and nuclear DNA release (21), which induced the hyperactivation of AIM2-ZBP1-ASC-RIPK1-RIPK3-CASP1-CASP8-GSDMD-GSDME (27), leading to cell membrane damage, and inducing membrane repair system (28), therefore, it is insensitive to the low concentration of cell membrane damaging agent Triton X-100, which leads to the production of resistance against T cell killing and immune evasion (22), but we can use IFN-γ and TNF-α to treat DLD1 and HCT116 to induce PANoptosis to overcome the immune escape of tumor cells; SW620 and HT29 (pMMR) do not have this mechanism, so they are sensitive to the low concentration of cell membrane damage agent Triton X-100, and they can be cleared by enhancing tumor immunity.

## Methods

### Cell culture

The human colorectal cancer cell lines DLD1, HCT116, RKO, SW480, SW0620 and mouse colorectal cancer cell lines CT26 (ATCC) were cultured in RPMI media (Corning, 10-040-CV) supplemented with 10% FBS and 1% penicillin and streptomycin. The human colorectal cancer cell lines HT29, embryonic kidney 293T cells and mouse MC38 were cultured in DMEM media (Corning, 10-040-CV) supplemented with 10% FBS and 1% penicillin and streptomycin.

### Plasmids

To knockout *Mlh1*, we used the gene editing crispr-cas9 system (lentiCRISPR-v2) (Addgene #52961). sgRNAs were designed using the CRISPR tool (http://crispr.mit.edu) to minimize potential off-target effects. Two sgRNAs, sgRNA1: TCACCGTGATCAGGGTGCCC and sgRNA2: ATTGGCAAGCATAAGCCATG to target mouse *Mlh1*, were cloned into lentiCRISPR-v2 plasmid. Human GSDMD-EGFP, GSDME-EGFP and mouse *Mlh1* lentiviral vector were purchased from (Igebio, China).

### Lentivirus production

Lentiviral particles were generated in HEK293T with co-transfection of above lentiCRISPR-v2, human GSDMD-EGFP, GSDME-EGFP or mouse *Mlh1* lentiviral vector and packaging plasmids pCMV-VSV-G (Addgene #8454) and psPAX2 (Addgene #12260) by Lipofectamine 3000 (Life technologies). Supernatant from HEK293T with transfection of Lentiviral packaging plasmids was collected, passed through a 0.22 μm filter to remove cell debris and frozen at −80 °C.

### Establishment of stable expression cell lines

For generating stable expression of human GSDMD-EGFP or GEDME-EGFP in DLD1 or HCT116, lentivirus particles were added to cell culture for 48h, infected cells were selected by 10 μg/ml Puromycin (MP Biomedicals) for two days.

### Gene editing

Mouse colorectal cancer cell line CT26 was infected with above *Mlh1* sgRNA-CRISPR-Cas9-expressing lentivirus at approximately 60% confluence in the presence of 5 μg/ml polybrene (Solarbio, China). 10 μg/ml Puromycin (MP Biomedicals) was used to select infected cells. Two days after selection for Puromycin, infected CT26 cells were single-cell cloned in 96-well plates by flow cytometry using the Beckman Coulter MoFlo XDP cell sorter. *Mlh1*-knockout clone was verified by western blot and DNA sequencing.

### Cell stimulation

For induction of cell death, 50 ng/mL of TNF-α (Peprotech, AF-300-01A), 100 ng/mL of IFN-γ (Peprotech, 300-02), 0.01, 0.008, 0.006, 0.004, 0.002% or 0.001 % v/V of Triton X-100 was used for the indicated time. For the inhibition of cell death, cells were co-treated with 20 μM of IDN-6556 (Sellck), 20 μM of Z-IETD-FMK (Sellck), 50μM of Nec-1 (Sellck), 20 μM of Necrosulfonamide (Sellck).

### Cytotoxicity assay

Relevant cells were treated as indicated. Cell viability was determined by the Cell Counting Kit-8 (Dojindo Laboratories). Cell death was measured by the LDH assay using CytoTox 96 Non-Radioactive Cytotoxicity Assay kit (Promega).

### Real-time imaging for cell death

The kinetics of cell death were determined using the ImageXpress Pico (Molecular Devices) live-cell automated system. Cells (2 X 10^3^ cells/well) were seeded in 96-well tissue culture plates. Cells were treated with the indicated cytokines and inhibitor inhibitor, then stained with propidium iodide (Sigma). The plate was scanned, and fluorescent and phase-contrast images (5 image fields/well) were acquired in real-time every 1 h from 0 to 96 h post-treatment. The images were analysed using the software package supplied with the ImageXpress Pico.

### Subcellular fractionation

According to the manufacturer’s instructions (Abcam), cells were seeded in 100 mm tissue culture plates and grown to semi-confluent density. Cells were detached by treatment with 2 ml of 0.25% Trypsin-EDTA and added into the saved medium. Cells were collected cells by centrifugation for 5 min at 300 x g at RT. Cell pellets were re-suspended in 1X Buffer A to 6.6 x 106 cells/ml. The equal volume of Buffer B was added to the cell suspensions and mixed by pipetting. Samples were incubated for 7 minutes on a rotator at RT and centrifuged at 5,000 x g for 1 min at 4°C. All supernatants were carefully removed and re-centrifuged at 10,000 x g for 1 min for the cytosolic fractions (C). The sequential cytoplasm-depleted cell pellets were re-suspended and combined in 1X Buffer A. Exactly the same volume of Buffer C were added to the suspensions and mixed by pipetting. Samples were Incubated for 10 minutes on a rotator at RT. Samples were centrifuged at 5,000 x g for 1 min at 4°C. All supernatants were carefully removed and re-centrifuge the supernatant fractions at 10,000 x g for 1 min for the mitochondrial fractions (M). The sequential cell pellets were re-suspended and combined in 1X Buffer A to the original volume of suspensions for the nuclear fractions (N). Samples were prepared by adding an appropriate amount of 5X SDS loading buffer and heated to 95 °C for 10 min for immunoblotting.

### Immunoprecipitation

Cells were lysed in a RIPA buffer containing (50mM Tris(pH7.4), 150mM NaCl, 1% NP-40, 0.25% sodium deoxycholate, protease inhibitors (Fudebio), phosphatase inhibitors (Fudebio)). After centrifugation at 20,000*g* for 10 min, the supernatant were incubated with either IgG control antibody (sc-3877, Santacruz) or anti-ASC antibody (67494-1-Ig, Proteintech) with protein A/G PLUS-Agarose (Santacruz) overnight at 4 °C. After washing with the above lysis buffer, the immunoprecipitated proteins were collected by centrifugation at 20,000*g* for 10 min, resuspended and boiled in 1× SDS loading buffer at 95 °C for 10 min.

### Immunoblot analysis

For immunoblot analysis, cells were lysed in RIPA buffer (containing protease inhibitors (Fudebio), phosphatase inhibitors (Fudebio)). Cells were centrifuged at 20,000*g* for 15 min at 4 °C, the soluble fraction was collected. Protein concentrations of cell lysates were determined using a BCA Protein Assay Kit (BeyotimeBio). Samples were prepared by adding an appropriate amount of 5X SDS loading buffer and heated to 95 °C for 10 min for immunoblotting. Proteins were separated by electrophoresis through 12% polyacrylamide gels. Following electrophoretic transfer of proteins onto PVDF membranes (Roche), nonspecific binding was blocked by incubation with 5% BSA, then membranes were incubated with primary antibodies against: caspase-3 (9662, Cell Signaling Technology (CST)), cleaved caspase-3 (9661, CST), caspase-7 (9492, CST), cleaved caspase-7 (9491, CST), caspase-8 (9746, 4790, CST), cleaved caspase-8 (8592, CST), caspase-9 (9504, CST), caspase-1 (ab179515, Abcam), GAPDH (60004-1-Ig, Proteintech), pMLKL (37333, CST, ab187091, Abcam), tMLKL (26539, CST), pRIPK1 (31122, CST), tRIPK1 (3493, CST), GSDMD (ab209845, Abcam, 39754, CST), GSDME (19859, Abcam), b-actin (66009-1-IG, Proteintech). Membranes were then washed and incubated with the appropriate horseradish peroxidase (HRP)–conjugated secondary antibodies (CoWin Biotech, anti-rabbit (CW0103) 1:10000, anti-mouse (CW0102) 1:10000. Proteins were visualized using Immobilon Forte Western HRP Substrate (Millipore, WBLUF0500).

### Immunofluorescence staining

In brief, cells were fixed in 4% paraformaldehyde (Yongjinbio) for 10 min and permeabilized with PBST containing 0.5% Triton X-100 for 10 min. Cells were then incubated in PBST containing 1% BSA for 30 min. Cells were incubated in primary antibodies (cleaved N-terminal GSDMD (ab215203, Abcam), cleaved N-terminal GSDME (ab222408, Abcam), pMLKL (ab187091, Abcam) overnight at 4 °C in PBST with 1% BSA. Cells were washed three times in PBST, then incubated in the appropriate secondary antibodies (Abcam) for 1 h at room temperature. Cells were washed three times in PBST. Cells were counterstained with DAPI (S2110, Solarbio). Images were acquired by confocal laser scanning microscopy (LSM780; Carl Zeiss) using 40× Apochromat objective.

## Author contributions

J.M.L. conceived the study; H.Y.L. performed all the experiments. H.L.N., Y.L., L.H.C and A.J.Z. provided technical assistance. X.K.Q., Y.Q.L., H.M. and C.Q. prepared the samples and materials. H.Y.L and J.M.L. wrote the manuscript.

## Acknowledgements

This work was supported by the National Key R&D Program of China (2017YFC1309000), the National Natural Science Foundation of China (81525020 and U1801282), the Guangzhou Science and Technology Plan Projects (Health Medical Collaborative Innovation Program of Guangzhou) (201803040019), the Guangdong Province Key R&D Program (2019B020229002).

